# Temperature impacts on mating and oviposition of the emerald ash borer, *Agrilus planipennis* Fairmaire (Coleoptera: Buprestidae)

**DOI:** 10.64898/2026.01.27.701830

**Authors:** Kenneth W. Dearborn, Daegan J. G. Inward, Sandy M. Smith, Chris J. K. MacQuarrie

**Affiliations:** Graduate Department of Forestry, John H. Daniels Faculty of Architecture, Landscape and Design, University of Toronto, Toronto M5S 3B3, Ontario, Canada; Canadian Forest Service, Natural Resources Canada, Great Lakes Forestry Centre, Sault Ste Marie, Ontario P6A 2E5, Canada; Forest Research, Alice Holt Lodge, Farnham, Surrey GU10 4LH, U.K

**Keywords:** invasive species, reproductive biology, thermal effects, fecundity, survival, climate

## Abstract

Local temperatures can shape the ability of introduced species to flourish and disrupt novel environments. The emerald ash borer (EAB), *Agrilus planipennis* Fairmaire (Coleoptera: Buprestidae), is an invasive beetle that threatens ash trees in North America and Europe. To assess the role of temperature on EAB reproduction, we reared groups of adult beetles at one of four temperatures (12, 15, 18, and 21 °C) and measured reproductive success (laying fertilized eggs and egg hatching). There was no effect of rearing temperature on EAB female lifespans but no eggs laid at 15 or 18 °C hatched, suggesting these temperatures disrupt the reproductive process of EAB. Females reared at 21 °C, however, consistently laid eggs that hatched. We then used these results to assess the likelihood of reproductive success over the previous ten years in eight cities in Canada that host EAB. All locations experienced temperatures of ≥ 21 °C, but the number of hours and the number of days above this critical temperature were highly variable. There were ample opportunities in all locations for EAB to reproduce, but EAB in cooler cities would experience thermal limitations thus slowing the spread of EAB populations.

## Introduction

Temperature regulates insect growth and development (Ludwig, 1928) and reproductive success (Zverev *et al*., 2018; Baur *et al*., 2022; Pilakouta & Baillet, 2022). Therefore, invasions by insects are climatically filtered, with local temperatures dictating how well introduced species can disrupt ecosystems. This climatic filtering acts as a pressure that can strain or disrupt key behaviours and developmental steps that insects must complete during their life cycles and is thus likely an important factor in why most introduced species fail to become pests (Williamson & Fitter, 1996). For example, temperature can alter and limit insect reproduction via effects on behaviour (Sih *et al*., 2002; Caetano *et al*., 2017; Augustin *et al*., 2022; Baur *et al*., 2022), physiology (Taylor, 1963; Niziolek *et al*., 2013), movement (Vasudeva *et al*., 2018; Zverev *et al*., 2018; Augustin *et al*., 2022; Sih *et al*., 2002), the efficacy of sperm transfer (Suzaki *et al*., 2018), and oviposition (Goebel, 2006; Keena *et al*., 2021). Disfunction caused by extreme weather can also create short- and long-term barriers, for example by preventing some or all of a population from being able to successfully mate (Vasudeva *et al*., 2018; Augustin *et al*., 2022; Sih *et al*., 2002) and overall climate may completely exclude an otherwise successful invasive species from expanding to certain spaces, despite appearing to have all the necessary physical resources (Goebel, 2006; Formby *et al*., 2018). The compounding effects of weather and climate can therefore challenge our ability to predict where in an invaded landscape an invasive species could be climatically limited.

The emerald ash borer (EAB), *Agrilus planipennis* Fairmaire (Coleoptera: Buprestidae), is a wood boring beetle that kills ash (*Fraxinus* spp.) (Oleaceae) trees in North America (Cappaert *et al*., 2005; CFIA, 2024; USDA, 2024) and elsewhere outside its native range (Orlova-Bienkowskaja, 2014). This species exploited an open niche in the phloem and cambium of North American ash trees (Gandhi & Herms, 2010; Paiero *et al*., 2012) and rapidly expanded its invasive range. Emerald ash borer likely co-evolved with Manchurian ash, *F. mandshurica* Ruprecht, in northeast Asia (Liu *et al*., 2003; Wei *et al*., 2004) and has successfully colonized the “naïve” ash species of North America (Cappaert *et al*., 2005) and Europe (Orlova-Bienkowskaja, 2014). At high larval densities, ash are overwhelmed and killed in as few as 2–3 years (Wang *et al*., 2010). The result is that hundreds of millions of ash trees have been killed by EAB larvae (Cappaert *et al*., 2005; Poland & McCullough, 2006).

The combined effects of climate and temperature have played an important role in the successful invasion of North America by EAB. Emerald ash borer spend most of their lives as larvae beneath the bark. This concealed existence allows populations to grow undetected and protects them, somewhat, from weather extremes (Vermunt *et al*., 2012; Duell *et al*., 2022). Adult EAB that live outside of trees, however, experience ambient air temperatures that impact their physiological processes (e.g., Fahrner *et al*., 2015). In North America the insect has invaded across a wide latitudinal distribution, from as far south as McLennan County, Texas (31.5° N) and north to Winnipeg, Manitoba (49.9° N) (CFIA, 2024; USDA, 2024). This vast distribution across North America means that adult emergence spans from as early as March in the south to as late as June in the north (MacDonald *et al*., 2022; Barker *et al*., 2023). This consequently means oviposition starts substantially later in the north (Barker *et al*., 2023). One consequence of this delay is that populations that inhabit the northern parts of EAB’s native and invasive ranges have a greater frequency of two-year life cycles (Wei *et al*., 2007; Tluczek *et al*., 2011; Jones *et al*., 2020), due to the cooler climates extending larval development (Duan *et al*., 2013).

The northward expansion of EAB in North America introduces temperature related uncertainties about EAB’s biology. Adult EAB must feed on ash foliage to complete development and become sexually mature (Pureswaran & Poland, 2009). This process takes up to 20 days for females when reared at constant temperatures of 20–25 °C (Rutledge & Keena, 2012; GLFC IPS, 2018). During the adult stage, cooler temperatures also impact EAB behaviours and physiology which could curtail mate finding and ultimately reduce reproductive success (Fahrner *et al*., 2015; Lelito *et al*., 2007). Range expansion of other invasive wood boring insects into regions with cooler temperatures occurs more slowly compared to range expansion of wood boring insects into regions with warmer temperatures, such as the tropics (Lantschner *et al*., 2014, Straw *et al*., 2016, Ward & Riggins, 2023). Temperatures of 25–28 °C are the optimal range for EAB mating and egg production (Duan *et al*., 2013; Hoban *et al*., 2016; GLFC IPS, 2018) but data are not available for lower temperatures more typical of climate and weather in the insect’s northern invaded range. Under natural conditions with varying temperatures, we would therefore predict that the maturation, and subsequent oviposition, periods are extended by cooler temperatures. If so, this likely has cascading effects on EAB development, leading to delayed egg hatch and larval development (Duan *et al*., 2013) and subsequent impacts on dispersal.

We investigated the role of temperature on the mating-fertilization-oviposition component of EAB’s reproduction pathway to develop predictions about how invasive spread of EAB could be influenced by these life history parameters. Since EAB populations at more northern latitudes experience cooler temperatures as adults, we hypothesized that this could contribute to slower spread and population growth. To test this hypothesis, we examined the role of cooler temperatures (12–21°C) on EAB reproduction. We then used a representative sample of major Canadian cities and their 10-year climate data to assess if those locations provided enough days at temperatures suitable for mating and oviposition.

## Methods

### Insects

We collected EAB adults from wild, infested green ash, *Fraxinus pennsylvanica* Marshall. All trees were harvested from forests in southern Ontario, Canada in the fall of 2021 and exhibited symptoms of infestation by EAB. These harvested trees were sectioned into approx. 30 cm lengths and transported to the Great Lakes Forestry Centre (GLFC) in Sault Ste. Marie, Ontario, Canada. We held the ash logs infested with EAB larvae outdoors at ambient winter temperatures in an unheated sea container (Fick & MacQuarrie, 2018; Dearborn *et al*., 2025) from fall 2021 until March 2022 when they were moved into a 4 °C environment chamber. The length of storage time does not impact adult fitness (Fick & MacQuarrie, 2018).

In February 2022 and May of 2022, we removed sections of ash from either the sea container or the environment chamber and placed them in emergence cages held at 27 °C, 60% relative humidity, and a 16:8h light:dark photoperiod to stimulate emergence of EAB adults. Under these conditions adults typically begin to emerge after 21 days (GLFC IPS, 2018). Once emergence began, we collected adults from the cages every 24 hours during morning light hours. We determined the sex of all newly emerged beetles by the presence (males), or absence (females), of gold-coloured setae on the ventral thorax (Cappaert *et al*., 2005; Rodriguez-Soana *et al*., 2007; Paiero *et al*., 2012). Adult EAB require a period of maturation feeding before they become receptive to mating and before they can produce viable eggs (Pureswaran & Poland, 2009). Males and females were kept separate for 14 days to ensure they had become sexually mature before combining males and females for mating (GLFC IPS, 2018; Dearborn *et al*., 2025). For this period of maturation feeding, we placed up to 8 adult beetles in single-sex groups in 946 mL (32 oz.) cups with perforated plastic lids (Placon; Madison, Wisconsin), hereafter referred to as feeding groups. Each feeding group consisted of male or female adults that were collected from emergence cages on the same day, and so were all considered to be the same age.

We maintained all feeding groups in a walk-in environment chamber (Hotpack Model 8112; Waterloo, Ontario) at 25 °C, 55% RH, and 16:8h light:dark photoperiod. We maintained all EAB at 25 °C to ensure they reached sexual maturity (GLFC IPS, 2018). We provided foliage from tropical ash, *Fraxinus uhdei* (Wenzig) Lingelsheim, from a greenhouse population maintained at the GLFC as a food source for each feeding group. We kept the foliage in good condition by inserting the stems through a hole in the lid of a scintillation vial (20 ml) that contained water.

We removed all dead EAB during daily inspections and moved feeding groups to clean cups and provided fresh foliage every three or four days. The maturation feeding period lasted 14 days from the time the feeding group was created.

### Mating Experiments

We evaluated the effect of temperature on mating and egg production using small groups of males and females. Hereafter we refer to these as mating groups. After the 14 days of sexual maturation within their assigned feeding groups, we created up to three male-female pairs and placed them into a fresh cup with foliage and water, as described above. All females within a mating group were selected such that they emerged on the same day. We did this to allow us to make an accurate determination of female lifespan. Cups housing the mating groups were the same as those used in the 14-day maturation period, but the plastic lid was replaced with a mesh screen covered by a coffee filter paper held in place with elastic bands. This coffee filter-mesh combination acts as a suitable substrate for female EAB to lay eggs (Dearborn *et al*., 2025).

We assigned the mating groups to one of four temperature treatments: 12, 15, 18, and 21 °C. We held all mating groups in reach-in environmental chambers (Conviron CMP 3244, Controlled Environments LTD, Winnipeg, Manitoba) at 55 % relative humidity and a 16:8h light:dark photoperiod. Males and females within each mating groups died over the course of the experiment. If all the females in a mating group perished before the males, we redistributed those males to other mating groups within the same temperature treatment that were without males. We did this to ensure females had the opportunity to mate throughout the experiment.

We inspected the filter paper for each mating group for eggs every 24 hours during the light hours of the photoperiod. If eggs were present, that filter paper was removed and replaced with a new filter paper. The filter papers with eggs were then moved to a different environmental chamber (Hotpack Model 8112; Waterloo, Ontario) at 25 °C, 55% RH, and 16:8h light:dark photoperiod to develop through to hatching. Emerald ash borer eggs do not hatch at 12 °C (Duan *et al*., 2013) and 25 °C is an optimal temperature to elicit egg development (GLFC IPS, 2018). We then counted the total number of eggs oviposited and the total number of hatched eggs produced by each of the mating groups in each of the four mating temperature treatments. Freshly oviposited EAB eggs are creamy white and will darken to a brown colour if they are fertilized (Rutledge & Keena, 2012). Female EAB will also sometimes deposit unfertilized eggs, or eggs that fail to develop. Eggs that darkened to brown were deemed to be ‘viable’ eggs (i.e., with the potential to produce a larvae). Hatched eggs were distinguished by being clear and with either the presence of an emerged larvae near the egg, or a small hole in the eggshell created by the larvae’s mandibles.

We repeated this experiment twice, using identical methods except as described here. In our first experiment we did not observe any oviposition (i.e., no eggs were found on the filter paper) by the mating groups held at 15 °C (n = 12 mating groups) or 18 °C (n = 12) after 19 days from the initiation of adult mating group formation. We did, however, observe oviposition by the mating groups held at 12 °C (n = 13) and 21 °C (n = 12) and so the lack of oviposition by mating groups in the 15 °C and 18 °C treatments was unexpected. To investigate this behaviour, we moved a subset of five mating groups from each of the four treatment temperatures to 25 °C for 12 days. Female EAB consistently lay eggs at 25 °C (Dearborn *et al*., 2025) and so this was an attempt to elicit oviposition under optimal conditions. We subsequently observed that at least 80% of the relocated mating groups from all temperature cohorts oviposited at 25 °C. These hatched eggs confirmed fertility for each subset of beetles. Once we had made that determination the subsets of mating groups were viable, we returned them to their assigned mating temperatures and monitored until all the females died. In the second repetition of the experiment, we followed the methods as described above, however, we assigned the four temperatures to different environmental chambers from the first experiment. This was an attempt to reduce chamber effects on oviposition. The second experiment had eight mating groups held at 12 °C, seven for both 15 and 21 °C, and six for 18 °C.

### Reproductive Timing in Canadian Cities

To assess the temperatures EAB would experience during their reproductive activities across their northern range we obtained the hourly and daily weather data for June 1st through October 31st during the years of 2010–19 from the Government of Canada (ECCC, 2024) for eight Canadian cities with known EAB populations. We then extracted the daily maximum air temperature to assess how many days per year adult female EAB in each city would experience a temperature ≥ 21 °C (i.e, warm enough to elicit mating, egg lay, and egg hatch based on the previous section). We also used this data set to determine the average day of the year when EAB would experience their first day ≥ 21 °C in each city. Finally, using the same data set to determine the number of hours ≥ 21 °C each year that EAB would encounter in each location. We selected Winnipeg, Manitoba; Sault Ste. Marie, Ontario; Windsor, Ontario; Toronto, Ontario; Montreal, Quebec; Quebec City, Quebec; Fredericton, New Brunswick; and Halifax, Nova Scotia as representative cities because they span the known invaded range of emerald ash borer in Canada at the time of this study. Each city hosts multiple weather stations, so we selected data from those that were complete for the 10 years under consideration. We limited our consideration to the months of June through October because very few EAB females emerge before 1 June, and those that do are unlikely to complete sexual maturation before that date (Barker *et al*., 2023) and temperatures in Canada after October 31 are typically too cool to elicit emergence by the insect, and ash leaves have senesced and are no longer present on trees, or are of such low quality that adult feeding cannot be supported.

### Statistical Analysis

We assessed the effect of mating temperature on the lifespan of female EAB with a Kaplan-Meier survival analysis. Only females that survived > 14 days were included in this analysis, as those individuals that perished during the maturation feeding stage were not assigned a mating temperature. We used a generalized linear model (GLM) to test if the number of hours per year ≥ 21°C differed among the eight Canadian cities over the 10-year period of interest. Analyses were performed using the R statistical language (R Core Team, 2022) using the Survival package (v3.5-0; Therneau *et al*., 2023). All figures were produced using ggplot2 (v3.4.1, Wickham *et al*., 2023).

## Results

### Mating Experiments

In experiment 1, a total of 28 or 29 females were exposed to each treatment temperature. To be assigned to a temperature treatment and a mating group, beetles needed to survive at least 14 days. Average female lifespan (mean ± SD) was shortest in those mating groups at 21 °C, 41.8 ± 14.9 days, and longest in those at 18 °C, 65.9 ± 33.0 days (Table 1). The shortest-lived females were reared at 12 and 21 °C (16 days) while the longest-lived individual female (141 days) was reared at 15 °C. In experiment 2, 17 or 18 females were exposed to each treatment temperature. Mean female lifespan within a rearing temperature was lowest at 12 °C, 36.4 ± 32.2 days, and highest at 18 °C, 47.4 ± 34.7 days (Table 1). The shortest-lived females among all temperature treatments survived 16 days and the longest-lived individual female (134 days) was reared at 12 °C (Table 1). We found an effect of mating temperature on lifespan of adult EAB females in experiment 1 (χ^2^ = 10.3; df = 3; P = 0.02; Figure 1A) but not in experiment 2 (χ^2^ = 0.6; df = 3; P = 0.9; Figure 1B). In experiment 1, there was no effect of the 12-day temperature transfer on female lifespan (χ^2^ = 0.0; df = 1; P = 0.9). We then combined data from both experiments and repeated the analysis of lifespan. In that analysis we did not detect an effect of mating temperature on lifespan (χ^2^ =6.1; df = 3; P = 0.1; Figure 1C).

**Figure 1.**
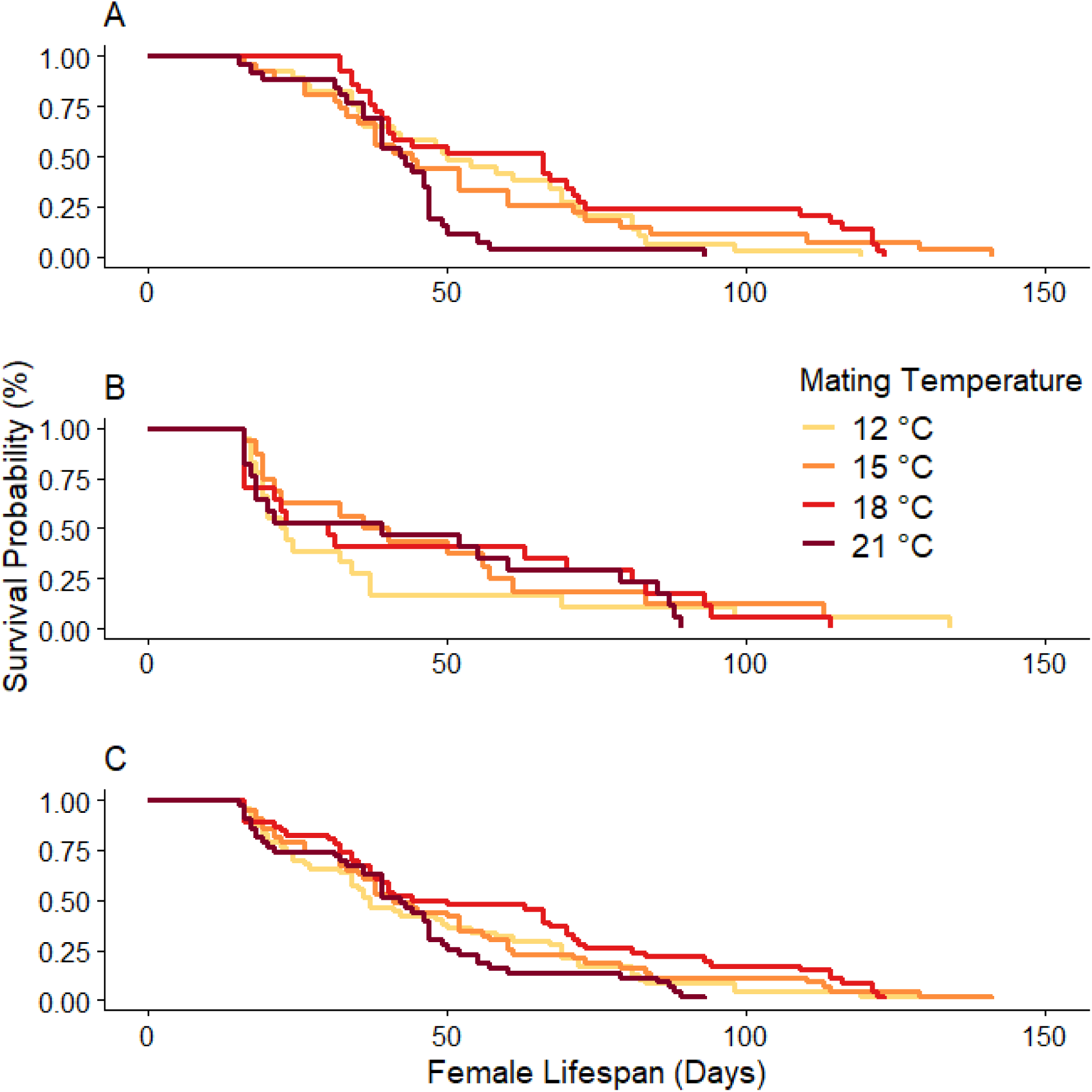
Probability of survival of adult female emerald ash borer (*Agrilus planipennis*) reared at one of four constant mating temperatures: 12 °C, 15 °C, 18 °C, or 21 °C in experiment 1 wherein a subset of insects were exposed to 12 days at 25 °C (A) or in experiment 2 where no insects were transferred (B), and the probability of survival when data from both experiments are combined (C).

**Table 1.**
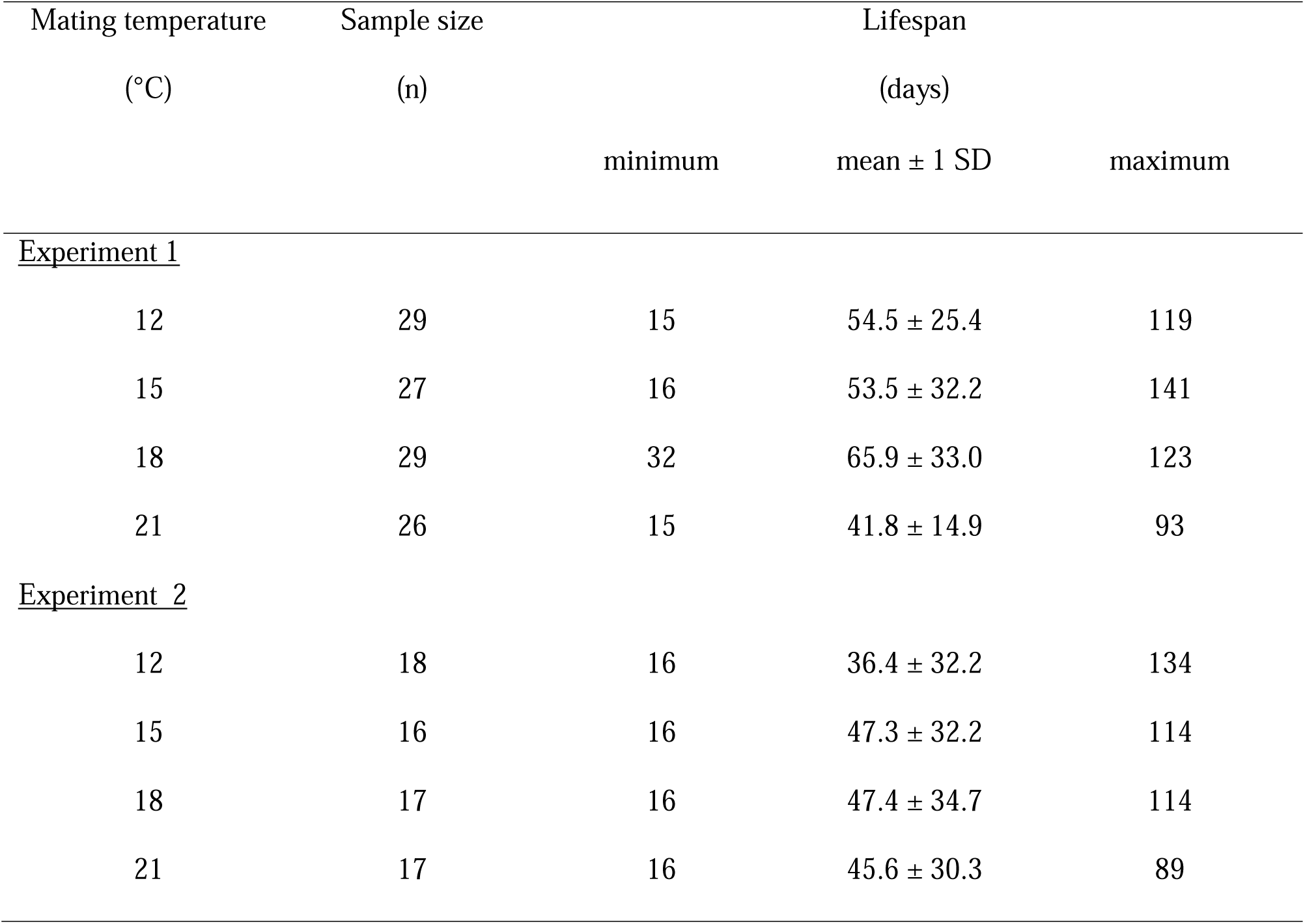
Lifespans of female emerald ash borer (*Agrilus planipennis*) reared at four constant temperatures in two experiments.

| Mating temperature | Sample size | Lifespan |  |  |
| --- | --- | --- | --- | --- |
| (°C) | (n) |  |  |  |
|  |  | minimum | mean ± 1 SD | maximum |
| <u>Experiment 1</u> |  |  |  |  |
| 12 | 29 | 15 | 54.5 ± 25.4 | 119 |
| 15 | 27 | 16 | 53.5 ± 32.2 | 141 |
| 18 | 29 | 32 | 65.9 ± 33.0 | 123 |
| 21 | 26 | 15 | 41.8 ± 14.9 | 93 |
| <u>Experiment 2</u> |  |  |  |  |
| 12 | 18 | 16 | 36.4 ± 32.2 | 134 |
| 15 | 16 | 16 | 47.3 ± 32.2 | 114 |
| 18 | 17 | 16 | 47.4 ± 34.7 | 114 |
| 21 | 17 | 16 | 45.6 ± 30.3 | 89 |

Female EAB produced eggs in all four temperature treatments in experiment 1 (Table 2; Table 3). In experiment 1, females in the 12 and 21 °C temperature treatments laid their first eggs on day 21 (i.e., 7 days after they were placed into mating groups). Females in the 15 and 18 °C treatments laid their first eggs on day 31, excluding any oviposition occurring during the 25 °C temperature transfer period. In experiment 1, the mean time to first oviposition was not longer at lower temperatures as expected. The mean time (± SD) to first oviposition from adult emergence was 25.4 ± 2.7 days for the mating groups held at 12 °C (n = 13) and 26.4 ± 4.9 days for insects reared at 21 °C (n = 12). The subsets of five mating groups in the 12 and 21 °C treatments that were assigned to the 12-day transfer to 25 °C had all produced eggs prior to being moved to the warmer temperature. By the end of experiment 1, five mating groups in the 15 °C treatment produced eggs. However, four of those five mating groups did so only after the brief exposure to 25 °C. In contrast, only one mating group reared at a constant 15 °C produced eggs, and then only on day 31 of the experiment. One other 15 °C mating group laid 11 eggs after the 12-day 25 °C exposure. Eight mating groups in the 18 °C treatment also produced eggs. The subset of five that were exposed briefly to 25°C all produced their first eggs while in the warmer growth chamber. The three groups that experienced a constant 18 °C laid their first eggs after 43.7 ± 17.8 days. In experiment 2, we only observed oviposition by EAB reared at the two warmest treatment temperatures of 18 (3 of 6 mating groups, 50%) and 21 °C (7 of 7, 100%).

**Table 2.** Percentage of emerald ash borer (*Agrilus planipennis*) mating groups that produced at least one viable egg when reared at one of four constant temperatures, with and without transfer to 25 °C during the mating period. A subset of five mating groups from each assigned temperature were returned to 25 °C for 12 days during the first experiment.

|  | Constant Rearing Temperature (°C) |  |  |  |  |  |  |  |
| --- | --- | --- | --- | --- | --- | --- | --- | --- |
|  | 12 |  | 15 |  | 18 |  | 21 |  |
| Transferred? | No | Yes | No | Yes | No | Yes | No | Yes |
| Experiment 1 | 76.9% | 80% | 0% | 80% | 16.7% | 100% | 91.7% | 100% |
| Experiment 2 | 0% | - | 0% | - | 16.7% | - | 100% | - |

**Table 3.** Number of eggs (mean ± 1 SD) produced by mating groups (n) of emerald ash borer (*Agrilus planipennis*) reared at one of four constant temperatures. Not all mating groups contributed eggs, * refers to the egg laying sample size if fewer than all mating groups oviposited.

|  | Constant rearing temperature (°C) |  |  |  |
| --- | --- | --- | --- | --- |
|  | 12 | 15 | 18 | 21 |
| Experiment 1 | 100.3 ± 69.4<br>(n = 13) | 7.0 ± 5.7<br>(n = 12, *2) | 30.2 ± 39.0<br>(n = 12, *6) | 82.0 ± 74.8<br>(n = 12) |
| Experiment 2 | 0<br>(n = 8) | 0<br>(n = 7) | 58.0 ± 34.0<br>(n = 6, *3) | 241.7 ± 161.9<br>(n = 7) |

In both experiments, viable eggs (i.e., those that darkened and should produce a larvae) were only produced by EAB females reared at 18 and 21 °C (Table 4). Most of these viable eggs, however, were produced by the mating groups reared at 21 °C. In experiment 2, we observed that 55.7% of eggs produced by females in the 21 °C were viable while just three viable eggs (1.7%) were produced by mating groups held at 18 °C (Table 4). EAB females reared at all other temperatures did not produce viable eggs. During the 12-day 25 °C rearing period all temperature groups produced more than 250 eggs (23.3–43.6 eggs/day) with viability and hatching percentages ranging from 47.3–73.7% and 41.9–60.2%, respectively. The subset of 5 matings groups moved to 25 °C from 15 and 18 °C reared EAB laid more eggs in 12 days than the full population did throughout experiment 1 exposed to the assigned cooler temperatures (i.e., 279 and 429 eggs compared to 14 and 181 eggs). The EAB mating groups reared at 12 and 21 °C in experiment 1 produced eggs that hatched. In experiment 2, however, only those eggs laid by EAB mating groups reared at 21 °C hatched (Table 4). None of the eggs that were laid by beetles reared at 15 or 18 °C hatched in either experiment (Table 4).

**Table 4.** Summary of reproductive success for emerald ash borer (*Agrilus planipennis*) reared at four constant temperatures in two experiments. ✓ refers to the event occurring and × is when there was not a reproduction event.

| Constant rearing temperature (°C) | Oviposited ? | Viable egg(s) laid ? | Egg(s) hatched ? | Fertility (% of viable eggs) | Hatch of total oviposition (%) | Hatch of viable eggs (%) |
| --- | --- | --- | --- | --- | --- | --- |
| <u>Experiment 1</u> |  |  |  |  |  |  |
| 12 | ✓ | ✓ | ✓ | 46.4 | 34.2 | 73.7 |
| 15 | ✓ | ✕ | ✕ | 0 | 0 | - |
| 18 | ✓ | ✓ | ✕ | 5.0 | 0 | 0 |
| 21 | ✓ | ✓ | ✓ | 60.5 | 43.0 | 71.1 |
| <u>Experiment 2</u> |  |  |  |  |  |  |
| 12 | ✕ | ✕ | ✕ | 0 | 0 | - |
| 15 | ✕ | ✕ | ✕ | 0 | 0 | - |
| 18 | ✓ | ✓ | ✕ | 1.7 | 0 | 0 |
| 21 | ✓ | ✓ | ✓ | 55.7 | 43.1 | 77.5 |

### Reproductive timing in Canadian Cities

We found a significant effect of city on hours ≥ 21°C (F(7, 72) = 63.298, p < 0.001). A post-hoc Tukey comparison of all pairs of cities found that 7/28 city pairs did not have significant differences in total hours ≥ 21°C. Winnipeg had significantly more total hours over 21°C than Halifax (p < 0.001) and Sault Ste. Marie (p < 0.001) despite being much further north. The eight Canadian cities all experience their first ≥ 21 °C day during the first week of June, on average (Table 5). Each city also experiences approximately the same average number of days with temperatures ≥ 21 °C (Table 5). Though cities further south (Windsor, Toronto, Montreal) average slightly more days above 21 °C between June and October than cities further to the north (Fredericton, Quebec City, Sault Ste Marie) (Table 5). Where we observed the most apparent differences among cities was in the total number of hours above 21 °C (Figure 2). Windsor had the greatest mean (± SD) number of total hours ≥ 21 °C from June through October at 1584.6 ± 188.0 hours, which was roughly triple what EAB in Halifax would experience at 519.3 ± 87.0 hours (Figure 2). Despite Winnipeg being by far the most northerly city investigated (49.9 ° N) the city experiences 1050.8 ± 142.6 total hours ≥ 21 °C, while the next three highest latitude cities (Quebec City, Sault Ste Marie, and Fredericton) all receive fewer than 850 total hours ≥ 21 °C (Figure 2). Halifax, is a coastal city which likely contributed to the low mean total hours ≥ 21 °C.

**Figure 2.**
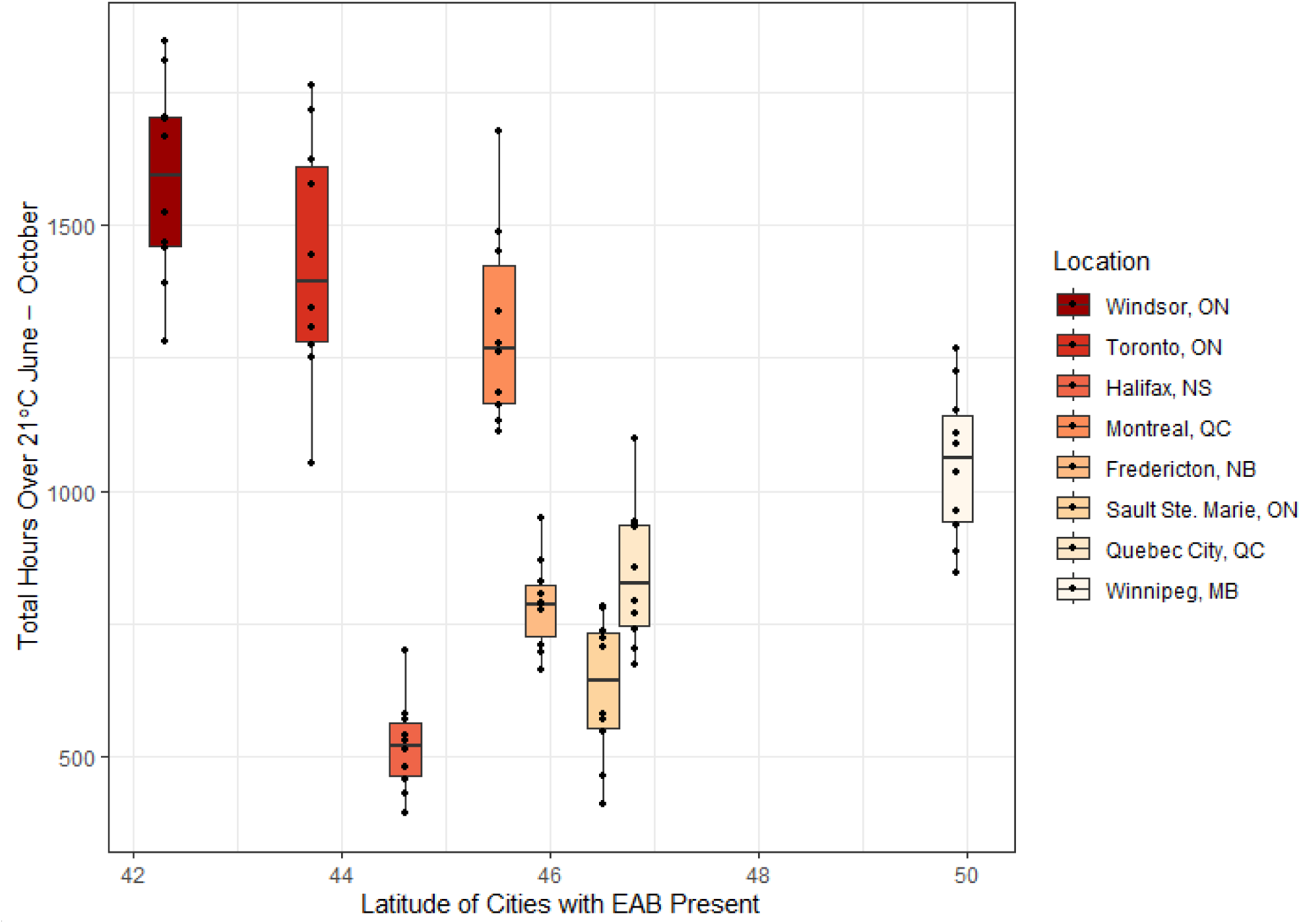
The total number of hours ≥ 21 °C between June 1^st^ and October 31^st^ in 2010–2019 for eight cities in Canada with established populations of emerald ash borer (*Agrilus planipennis*) (dots), ordered by latitude. Boxes show the mean (thick black line) and 25th and 75th percentiles, whiskers are 1.5 times the distance between the 1st and 3rd quartiles of the data.

**Table 5:** Number of days with temperatures > 21 °C and the mean first date when a temperature > 21 °C is observed for ten cities in Canada with established populations of emerald ash borer (*Agrilus planipennis*). Summaries were calculated from temperature data collected between June 1st and October 31st in 2010–2019. The average first day with a temperature of > 21°C was determined from the mean ordinal date and then converted to the calendar date.

| City<br>(weather station coordinates) | Number of days > 21 °C (mean<br>± SD) | Mean first date with a<br>temperature > 21 °C |
| --- | --- | --- |
| Fredericton, New Brunswick<br>(45.92° N, 66.61° W) | 89.5 ± 5.9 | June 2<sup>nd</sup> |
| Halifax, Nova Scotia<br>(44.63° N, 63.51° W) | 74.9 ± 4.2 | June 8<sup>th</sup> |
| Montreal, Quebec<br>(45.50° N, 73.58° W) | 98.4 ± 8.3 | June 4<sup>th</sup> |
| Quebec City, Quebec<br>(46.78° N, 71.29° W) | 85.7 ± 4.8 | June 3<sup>rd</sup> |
| Sault Ste. Marie, Ontario<br>(46.48° N, 84.51° W) | 82.1 ± 7.4 | June 3<sup>rd</sup> |
| Toronto, Ontario<br>(43.67° N, 79.40° W) | 103.4 ± 5.7 | June 2<sup>nd</sup> |
| Windsor, Ontario<br>(42.28° N, 82.96° W) | 109.7 ± 6.5 | June 1<sup>st</sup> |
| Winnipeg, Manitoba<br>(49.89° N, 97.13° W) | 92.6 ± 6.2 | June 2<sup>nd</sup> |

**Table 5:** Number of days with temperatures $> 21^{\circ}\text{C}$ and the mean first date when a temperature $> 21^{\circ}\text{C}$ is observed for ten cities in Canada with established populations of emerald ash borer (*Agrilus planipennis*).
| City<br>(weather station coordinates) | Number of days $> 21^{\circ}\text{C}$ (mean<br>± SD) | Mean first date with a<br>temperature $> 21^{\circ}\text{C}$ |
| --- | --- | --- |
| Fredericton, New Brunswick<br>(45.92° N, 66.61° W) | 89.5 ± 5.9 | June 2<sup>nd</sup> |
| Halifax, Nova Scotia<br>(44.63° N, 63.51° W) | 74.9 ± 4.2 | June 8<sup>th</sup> |
| Montreal, Quebec<br>(45.50° N, 73.58° W) | 98.4 ± 8.3 | June 4<sup>th</sup> |
| Quebec City, Quebec<br>(46.78° N, 71.29° W) | 85.7 ± 4.8 | June 3<sup>rd</sup> |
| Sault Ste. Marie, Ontario<br>(46.48° N, 84.51° W) | 82.1 ± 7.4 | June 3<sup>rd</sup> |
| Toronto, Ontario<br>(43.67° N, 79.40° W) | 103.4 ± 5.7 | June 2<sup>nd</sup> |
| Windsor, Ontario<br>(42.28° N, 82.96° W) | 109.7 ± 6.5 | June 1<sup>st</sup> |
| Winnipeg, Manitoba<br>(49.89° N, 97.13° W) | 92.6 ± 6.2 | June 2<sup>nd</sup> |

## Discussion

We found that EAB are capable of successfully mating and producing eggs at temperatures as low as 12 °C. Those eggs, however, are incapable of hatching if they are laid by mothers reared at moderate temperatures of 15 or 18 °C. In our second experiment, EAB females held at 21 °C were the only insects that produced eggs that hatched. This suggests temperatures < 21 °C inhibit the reproductive process and that 18 °C is near the lower thermal limit for either mating, fertilization, or oviposition. If so, then our results suggest that climate and weather limit the reproductive potential of EAB through direct effects on how long females are active and physiologically capable of producing viable eggs.

Local temperatures could affect the invasive pressure EAB exerts on North American ash trees. When the adult stage coincides with extended periods of 18 °C or lower our results suggest EAB can be active but not able to reproduce. The adult stage of EAB do not cause direct damage to trees (Wei *et al*., 2007) but drive subsequent larval feeding damage (Cappaert *et al*., 2005; Wei *et al*., 2007) by selecting oviposition sites (Rigsby *et al*., 2014). If relatively cool temperatures inhibit the insect’s reproductive pathway, then one consequence is that adults may die before reproducing. For example, avian predators - which are not hindered by cool weather - forage upon resting adult EAB (Flower *et al*., 2014). The combination of reduced reproductive capacity associated with cool temperatures and a consistent predation risk could slow local tree decline by decreasing overall adult reproductive success.

Extended EAB larval feeding at cooler temperatures provides a greater opportunity for biotic resistance from ash tree defenses, predation, and parasitism. Ash trees produce callus tissue that can slow larval development (Rutledge & Arango-Velez, 2017). Woodpecker feeding on larvae occurs year-round (Koenig *et al*., 2013) and is a significant EAB mortality factor (Koenig & Liebhold, 2017). Woodpeckers and other predators including parasitoids could help moderate the northward progression of EAB if climate slows the development of larvae (Duan *et al*., 2013; Barker *et al*., 2022) and increases the frequency of a two-year life cycle (Jones *et al*. 2020). It is generally assumed, however, that the lack of abiotic and biotic resistance across the invaded range of ash will allow for continued EAB spread. Our results suggest that this assumption is not supported and that abiotic conditions may limit the reproductive potential of EAB by limiting the amount of time available to mate at the northern extent of the insect’s range in North America and Europe.

Invasive populations of EAB will continue to expand their non-native ranges across North America and Europe. In all places where ash trees grow natively, there are multiple days warm enough to allow mating and reproduction to occur (Barker *et al*., 2023; Figure 2). The eight Canadian cities we looked at do not consistently experience temperatures that would inhibit the reproductive abilities of adult EAB. Populations of EAB in Windsor, Toronto, Montreal, and Winnipeg all receive > 1000 hours with temperatures ≥ 21 °C. This suggests there are sufficient opportunities for mating to occur. Halifax, despite having the fewest mean hours ≥ 21 °C of these cities, hosts a growing population of EAB. That there are fewer hours to mate, however, may explain the lower rate of female mating success in the Halifax population (Caouette *et al*., 2024). Emerald ash borer adults can also increase their internal temperatures behaviourally through solar irradiation (i.e., basking) as they are primarily active in the canopy (Lelito *et al*., 2007). Sun exposure can also increase temperatures on and under the bark (Vermunt *et al*., 2012) which benefits both egg and larval development, which is why female EAB select oviposition sites that are sunny, preferring the south-west facing side of ash trees (Timms *et al*., 2006). Our findings suggest these two behaviours are likely important factors in reproductive success, particularly in populations that experience fewer hours where temperatures are conducive for mating and oviposition of viable eggs.

Our results suggest our temperature manipulations disrupted EAB’s reproductive process. The EAB in our experiments produced eggs at all four temperatures but none of the eggs that were laid by females reared at 15 or 18 °C hatched. In experiment 1, however, we did observe that EAB reared at 12 °C produced eggs which is not consistent with the observed behaviour at 15 and 18 °C. This could have been a response to their brief exposure to a temperature of 22 °C every few days when we removed the cups from the environment chamber to provide new foliage. During this brief period the females in this treatment may have performed “egg dumping”, which is a behavioural response to maximise reproduction under variable thermal conditions (Aluja *et al*., 2011). We postulate that only the 12 °C females performed the egg dumping because the approximate 10 °C difference was large enough to elicit the stress related response. Experiment 2 had a smaller sample size (16–18 females) and fewer mating groups (6–8) than those of experiment 1 (26–29 females and 12–13 mating groups). We did not record handling time when replenishing the ash food source, but the length of time beetles were exposed to 22 °C would have been less during experiment 2 and may account for the discrepancy in oviposition among the insects reared at 12 °C. All aspects of insect biology have working and optimal ranges for activities. For example, the efficacy of sperm transfer can be reduced outside of critical temperatures (Suzaki *et al*., 2018; Vasudeva *et al*., 2018). We did not dissect female EAB to examine for the presence of a spermatophore and so we are not able to state if successful matings occurred at these lower temperatures. It is therefore possible that the lack of realized fecundity was because the EAB in our experiments were not able to complete their courtship and mating behaviours (Rutledge & Keena, 2019).

Emerald ash borer pose significant ecological and economic risks to uninvaded regions with ash trees. We argue the unexpected egg production in the 12 °C mating group in experiment 1 are valuable data despite us not observing the same effect in experiment 2. Emerald ash borer has demonstrated plasticity in their ability to tolerate extreme cold (Duell et al., 2022) and so could have similar plasticity in its thermal range for mating and reproduction. We suggest further experiments are required to understand these effects in EAB.

Insects naturally experience daily temperature variation. Exposing them to constant temperatures in climate-controlled environments may modify behaviour and/or lead to changes in physiological processes not seen in the wild. For example, oviposition and fecundity in insects reared at an optimal constant temperature can often be higher than that of insects reared at fluctuating temperatures (Singh *et al*., 2018). In our experiments, we investigated temperatures thought to be suboptimal for EAB development (Duan *et al*., 2013). It is likely that repeating our methods but rearing EAB at fluctuating temperatures could result in better reproductive performance than we found here for constant temperatures. We are not aware of any studies that have attempted to rear EAB at fluctuating temperature, although doing so might better emulate its potential performance under a changing climate or in its expanding range.

We found that rearing EAB at constant temperatures < 21 °C compromised the insect’s reproductive pathway. This suggests cooler climates and more northern latitudes could have abiotic resistance against the spread of EAB populations. Adult EAB that experience temperatures that delay or prolong mating and oviposition will oviposit eggs later in the season. The larvae that hatch from these eggs will emerge into an environment that slows their development (Duan *et al*., 2013). If oviposition is delayed too long, some EAB eggs may never accumulate enough degree days to hatch and if this phenomenon affected many eggs there would be subsequent impacts on larval density, and thus the rate and frequency of mortality of ash trees. Slower rates of tree death and lower rates of EAB spread will provide more time to manage and mitigate damage. These findings on the impact of temperature on adult reproduction will aid management decisions in more northern and cooler climate regions where the risks from EAB may differ from more southerly, warmer regions where this invasive beetle has thrived.

## Acknowledgements

We thank Misha Demidovich for maintenance of tropical ash in the greenhouse and Gene Jones for the collection of EAB-infested ash from southern Ontario. Financial support of Defra (UK) Future Proofing Plant Health, Natural Resources Canada, and scholarships from the University of Toronto to KWD are also gratefully acknowledged.

## Author Contributions

**KWD:** Conceptualization; Data curation; Formal analysis; Investigation; Methodology; Project administration, Resources, Software; Validation; Visualization; Writing – original draft; Writing – review & editing. **DJGI:** Funding acquisition; Supervision; Writing – review & editing. **SMS:** Funding acquisition; Supervision; Writing – review & editing. **CJKM:** Project administration, Resources; Software; Supervision; Validation; Writing – review & editing

## Conflicts of Interest

The authors declare no conflicts of interest.

## Data Availability Statement

The datasets are available from the corresponding author upon reasonable request.

## Notes

### Competing Interest Statement

The authors have declared no competing interest.

